# AutoPVS1: An automatic classification tool for PVS1 interpretation of null variants

**DOI:** 10.1101/720839

**Authors:** Jiale Xiang, Jiguang Peng, Zhiyu Peng

**Author notes:** These authors contributed equally to this work (Jiale Xiang, Jiguang Peng). Correspondence: Dr. Zhiyu Peng at BGI Park, No.21 Hongan 3rd Street, Yantian District, Shenzhen 518083, China.

## Abstract

Null variants are prevalent within human genome, and their accurate interpretation is critical for clinical management. In 2018, the ClinGen Sequence Variant Interpretation (SVI) Working Group refined the only criterion (PVS1) for pathogenicity in the American College of Medical Genetics and Genomics and the Association for Molecular Pathology (ACMG/AMP) guidelines. The refinement may improve interpretation consistency, but it also brings hurdles to biocurators because of the complicated workflows and multiple bioinformatics sources required. To address these issues, we developed an automatic classification tool called AutoPVS1 to streamline PVS1 interpretation. We assessed the performance of AutoPVS1 using 56 variants manually curated by ClinGen’s SVI Working Group and achieved an interpretation concordance of 95% (53/56). A further analysis of 28,586 putative loss-of-function variants by AutoPVS1 demonstrated that at least 27.6% of them do not reach a very strong strength level, with 17.4% based on variant-specific issues and 10.2% on disease mechanism considerations. Moreover, 40.7% (1,918/4,717) of splicing variants were assigned a decreased PVS1 strength level, significantly higher than frameshift and nonsense variants. Our results reinforce the necessity of considering variant-specific issues and disease mechanisms in variant interpretation, and demonstrate that AutoPVS1 is an accurate, reproducible, and reliable tool which facilitates PVS1 interpretation and will thus be of great importance to curators.

## Introduction

The decreasing cost of massively parallel next-generation sequencing (NGS) has made it widely used for clinical diagnosis of hereditary diseases in patients^1^ and preconception/prenatal expanded carrier screening in healthy couples.^2^ With hundreds and thousands of variants produced per individual by NGS-based tests, interpreting their clinical significance and associations with clinical observations presents a significant challenge. The range of difficulties includes coping with different isoforms of a transcript,^3^ determining variant frequency thresholds,^4^ appropriately interpreting the clinical context,^5^ and incorporating with continually updated knowledge.^6^ Moreover, subjectivity in weighing evidence and interpreting guidelines leads to a lack of consistency, with interpretations of the same variant varying substantially between laboratories.^7; 8^

To standardize variant interpretation, the American College of Medical Genetics and Genomics and the Association for Molecular Pathology (ACMG/AMP) published a joint consensus recommendation that provides a set of evidence-based guidelines.^9^ The only criterion in the guideline with a very strong strength level for pathogenicity is named PVS1; it was recommended for cases of a “null variant (nonsense, frameshift, canonical ±1 or 2 splice sites, initiation codon, single or multiexon deletion) in a gene where loss-of-function (LoF) is a known mechanism of disease”.^9^ The accurate interpretation of PVS1 is critical for variant interpretation because null variants are prevalent within the human genome (at least 100 per genome).^10^ Moreover, they are markedly enriched in low-frequency alleles,^11^ meeting the moderate evidence (PM2) criterion for pathogenicity in the ACMG/AMP guidelines.^9^ With the combination of PVS1_VeryStrong and PM2, clinical significance could easily reach a likely pathogenic classification, which would lead to actionable clinical interventions.

The ACMG/AMP guideline did not elaborate specific considerations for the use of the PVS1 criterion. To fill this gap, the Clinical Genome Resource (ClinGen) Sequence Variant Interpretation (SVI) Working Group refined PVS1 to include the following strength levels: very strong (PVS1_VeryStrong), strong (PVS1_Strong), moderate (PVS1_Moderate), or supporting (PVS1_Supporting).^12^ The refinements were mainly focused on issues specific to each variant type and considerations about disease mechanisms.^12^ Comprehensive curation of these specifications is expected to increase the consistency of PVS1 interpretation, but it also present hurdles to biocurators because of the complicated workflows and multiple bioinformatics sources required.

The ClinGen Expert Panels also proposed disease/gene-specific PVS1 recommendations, including germline *CDH1*,^13^ *PTEN*,^14^ and *MYH7*-associated inherited cardiomyopathies.^15^ For variants in these genes/diseases, PVS1 interpretation should follow the gene/disease-specific recommendations rather than the PVS1 evidence strength levels of ClinGen’s SVI Working Group, which are meant to provide general guidance across all diseases.^12^

Using computational tools to facilitate variant interpretation has proven feasible in several cases. CardioClassifier^16^ and CardioVAI^17^ are tools for variant classification in cardiovascular-related genes. Kleinberger and co-authors offered an openly-available, online tool to efficiently classify variants with the option to aggregate each variant into an exportable table.^18^ Li and Wang developed a semi-automatic tool named InterVar to generate predictions based on 18 criteria in the ACMG/AMP guidelines.^19^ To facilitate its use, they also developed a web server version, wIntervar. However, the web version cannot process some variant types, such as indels and splicing sites, significantly weakening the predictive power of InterVar for non-command-line users. While valuable, none of these tools incorporates the latest specifications for PVS1 interpretation.^12^

In this study, we developed AutoPVS1, an automatic computational tool to streamline PVS1 interpretations based on the recommendations outlined by ClinGen’s SVI Working Group. To make the tool broadly accessible, we also offer a graphical user interface to AutoPVS1 for users who are not familiar with the command line. Our results demonstrate that AutoPVS1 generates fast, reproducible and reliable PVS1 interpretation results.

## Material and Methods

Consistent with the recommendations of the ClinGen’s SVI Working Group,^12^ AutoPVS1 consists of two steps: 1) application of PVS1 criteria specific to each variant type and 2) disease mechanism considerations (Figure 1).

**Figure 1.**
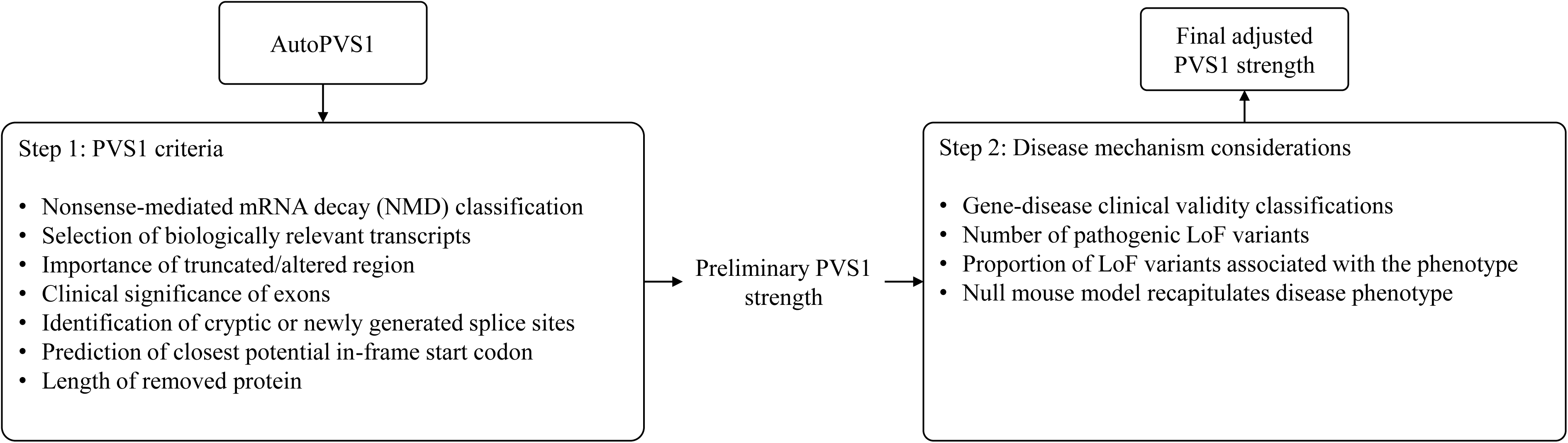
Illustration of the Two-Step Procedure of AutoPVS1.

### PVS1 criteria

The key criteria for PVS1 are nonsense-mediated mRNA decay (NMD) classification, selection of biologically relevant transcripts, the importance of the truncated/altered region, the clinical significance of exons, identification of cryptic or newly generated splice sites, and prediction of the closest potential in-frame start codon (Figure 1).

#### NMD classification

The NMD surveillance pathway is a mechanism that selectively degrades nonsense mRNAs which has been observed in all eukaryotes to date.^20^ In AutoPVS1, we determined that the NMD mechanism is active when a premature termination codon occurs in the 3’-most exon or within the 50 3’-most nucleotides of the penultimate exon.

#### Selection of biologically relevant transcripts

Transcript analysis plays a critical role in variant interpretation.^21^ Inappropriate transcript annotation may lead to missed or incorrect diagnoses.^3^ To annotate the biologically relevant transcript, we first retrieved the most prevalent transcripts used in ClinVar.^22^ The ClinVar 20190624 release was used throughout this study. If no result was found in ClinVar, we retrieved transcripts from RefSeqGene/the Locus Reference Genomic collaborations (see Web Resources). If both retrievals were unsuccessful, we used the longest available transcript in the Reference Sequence (RefSeq) database.^23^

#### Importance of the truncated/altered region

We extracted universal protein domain annotations from the University of California Santa Cruz (UCSC) Genome Browser Database.^24; 25^ We then retrieved missense variants with Pathogenic (P)/Likely Pathogenic (LP)/Benign (B)/Likely Benign (LB) classifications from ClinVar.^22^ Those with conflicting interpretations were excluded. To identify important truncated/altered regions, we considered only domains lacking B/LB variants and with at least five P/LP variants. If B/LB and P/LP variants were detected in the same region, we conservatively required that the number of P/LP variants be at least ten times greater than the number of B/LB variants. In addition, we included a list of functional regions curated by disease-specific experts (Table S1).

#### Clinical significance of exons

Evaluating the clinical significance of exon(s) is an essential step in determining how well a null variant meets the PVS1 criteria. The assessment includes two aspects: 1) the existence of the exon(s) in biologically relevant transcript(s); 2) the frequency of LoF variants in this exon in the general population. We used the transcript selected above as the most biologically relevant transcript(s) to assess the existence of exon(s).

To score the allele frequency, we retrieved data from the gnomAD database (v2.1) and filtered out low-quality variants.^26^ Given that a high frequency of LoF variants in the general population supports the clinical insignificance of an exon,^21^ we determined that exons were less important when their LoF variant(s) had an allele frequency greater than 0.1% in at least one gnomAD ancestry. Users should note that the allele frequency threshold can vary for different diseases.

#### Identification of cryptic or newly generated splice sites

To identify cryptic or newly generated splice sites, we assessed the nearby (±20 bp) sequences. If a potential splice site was not found, we extended the search to ±50 nucleotides.^27^ We applied MaxEntScan to predict the new site’s effect on splicing motifs.^28^ We used a MaxEntScan score above 3 or a >70% score of the canonical ±1,2 splice sites to identify splice sites (donor or acceptor), consistent with the threshold in the Human Splicing Finder.^29^

#### Prediction of the closest potential in-frame start codon

Given the evidence that alternative initial codons can be robust, ClinGen’s SVI Working Group suggested that no PVS1 strength level higher than moderate should be assigned to mutations in the initial codon.^12^ However, initial codon variants evaluated by ClinGen gene-disease specific Expert Panels had their PVS1 strength level elevated from moderate to very strong if they had ≥1 pathogenic variant(s) upstream of the closest potential start codon.^12^

We used a conservative threshold to determine the PVS1 strength level of initiation codon variants. We first extracted all 754 initiation codon variants from ClinVar and grouped them by their clinical significance. Next, we counted the number of P/LP variants upstream of the closest potential start codon for each initiation codon variant. The median number of each group was 7, 6, 0, 2.5 and 0 for P, LP, VUS, LB, and B initiation codon variants, respectively (Figure S1). We therefore determined the PVS1 strength level as follows: number of upstream variants >6 (PVS1_VeryStrong), 3 < number ≤ 6 (PVS1_Strong), 1 ≤ number ≤ 3 (PVS1_Moderate), number = 0 (PVS1_Supporting).

### Disease mechanism considerations

After assigning preliminary PVS1 strength levels via the PVS1 criteria, the second step in interpretation is to consider the disease mechanisms (Figure 1). ClinGen’s SVI Working Group recommends considering the number of pathogenic LoF variants, the proportion of LoF variants associated with the phenotype, the recapitulation of disease phenotype in null mice, and gene-disease clinical validity classifications in weighing a gene’s disease mechanism.^12^ We extracted the number of LoF variants from ClinVar. To assess the recapitulation of disease phenotypes in mice, we downloaded genes that have complete mouse models from the International Mouse Phenotyping Consortium.^30; 31^

With respect to gene-disease clinical validity, ClinGen developed an evidence-based framework to assess the strength of gene-disease relationships in six classifications: “Definitive”, “Strong”, “Moderate”, “Limited”, “No Reported Evidence,” or “Conflicting Evidence”.^32^ We aggregated genes and diseases curated by ClinGen’s experts in our analytic pipeline (see Web Resources). For those not yet curated, biocurators should manually adjust the strength levels.

### Gene/disease-specific PVS1 recommendations

Three gene-specific PVS1 criteria recommended by ClinGen Expert Panels that differ from the recommendations of ClinGen’s SVI Working Group are also included in our analytic pipeline. First, the ClinGen CDH1 Expert Panel recommended that PVS1_VeryStrong should be used for null variants, with the exception of canonical splice sites. Splicing variants should be assigned strong or moderate levels. Moreover, a moderate strength level (rather than a strong level) should be assigned if a premature stop is downstream of NM_004360.3:c.2506G>T (p.Glu836Ter).^13^ Second, the ClinGen PTEN Expert Panel recommended c.1121 position within exon 9 (NM_000314.4) as the 3’ boundary for PVS1_VeryStrong.^14^ Third, the ClinGen Inherited Cardiomyopathy Expert Panel assigned a moderate weight (PVS1_Moderate) to an LoF variant in the *MYH7* gene and removed the PVS1_VeryStrong rating.^15^

### Benchmarking

We constructed two datasets from the following sources. Dataset 1 included 56 variants of varying type that have been expertly curated by ClinGen’s SVI Working Group.^12^ To test the performance of AutoPVS1, we compared the PVS1 interpretation results of dataset 1 by ClinGen’s SVI working group, AutoPVS1, and InterVar^19^ (a generic tool for variant interpretation).

Since the second step of PVS1 interpretation requires gene-disease clinical validity, we constructed Dataset 2 using ClinVar putative LoF variants in 578 genes. Specifically, 577 genes were included because the gene-disease associations have been curated by ClinGen’s experts and are freely available (https://search.clinicalgenome.org/kb/gene-validity). One gene (*PAH*) has not been curated but was added to the list because of mutations which cause a common disease named Phenylketonuria (MIM: 261600). We manually curated the gene-disease association according to ClinGen’s framework^32^ and determined the *PAH*-Phenylketonuria association as “Definitive”. We excluded LoF variants in these 578 genes with low-quality calls in gnomAD database.^26^

We used Dataset 2 to assess the impact of PVS1 recommendations on variant interpretation. Since PVS1 interpretation consists of two procedures (Figure 1), we performed the assessment sequentially. We first assessed the PVS1 criteria to assign preliminary PVS1 strength levels. We then integrated disease mechanism considerations to assign adjusted PVS1 strength levels. In cases where a gene associated with more than one disease was curated and different gene-disease associations were determined, we conservatively chose the highest association levels for LoF variants in this gene in order to not exaggerate the ratio of decreased strength levels.

### A web interface for AutoPVS1

AutoPVS1 is a command-line program written in Python. Uploaded variants are annotated via Variant Effect Predictor.^33^ To assist users without command-line experience, we also offer a graphical user interface for AutoPVS1 (see Web Resources). Users can search for a null variant using sequence variant nomenclature recommended by the Human Genome Variation Society (HGVS)^34^ or chromosomal position. For exon-level deletions/duplications, the chromosomal initial and end positions of the exon(s) are required. When a variant is queried, AutoPVS1 first processes the PVS1 criteria to assign a preliminary PVS1 strength level and then initiates disease mechanism considerations, which are user-adjustable. We have

## Results

### High concordance with manually curated variants

To assess the performance of AutoPVS1, we used it to analyze all null variants (n=56) curated by ClinGen’ SVI Working Group, including 13 nonsense variants, 17 frameshift variants, 16 canonical ±1,2 splice variants, 4 initiation codon variants and 6 duplications/deletions.^12^ The interpretation consistency between AutoPVS1 and ClinGen’s SVI Working Group was 95% (53/56). One initial codon variant and two canonical ±1,2 splice variants were interpreted differently (Figure 2). We review the details of these three variants below.

**Figure 2.**
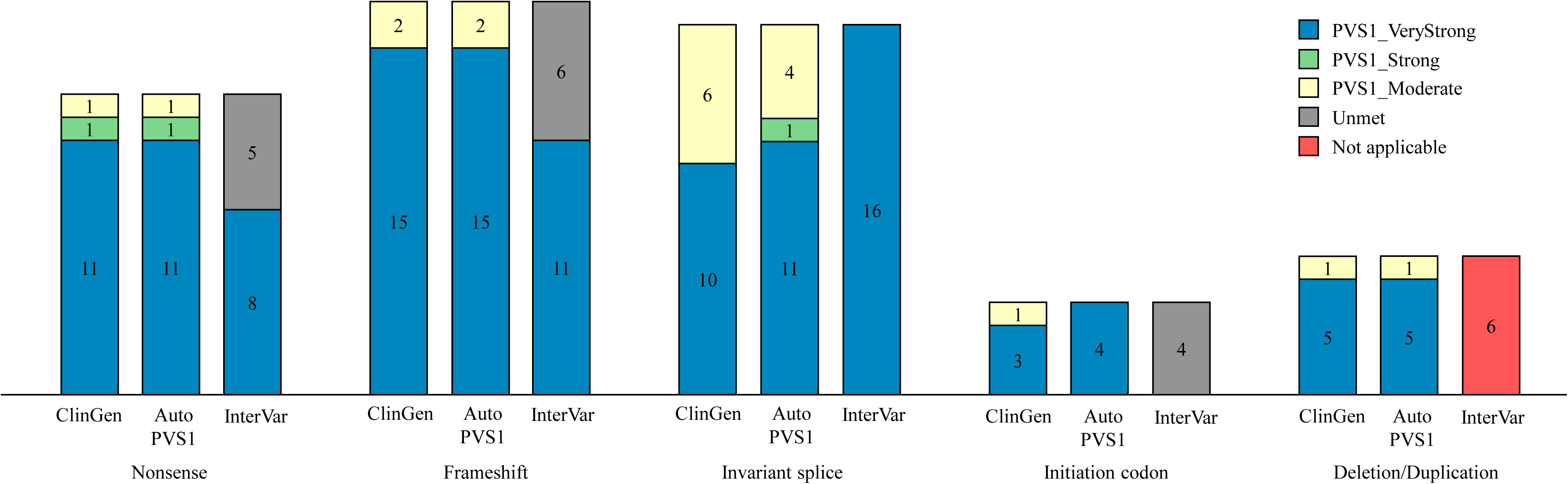
Comparison of AutoPVS1 with manually curated variants and InterVar. A total of 56 variants, of varying type, were used for comparison. There variants were manually curated by ClinGen Sequence Variant Interpretation Working Group. PVS1: the only criterion designated with very strong strength level for pathogenicity in the American College of Medical Genetics and Genomics and the Association for Molecular Pathology guideline. “Unmet”: None of strength levels was assigned. “Not applicable”: InterVar is not compatible with exon-level deletions/duplications.

Variant NM_000218.2:c.2T>C(p.Met1?) in the gene *KCNQ1* was rated PVS1_Moderate in ClinGen’s SVI Working Group’s classification, whereas it was classified as PVS1_Very Strong by AutoPVS1. The next methionine is at codon 159 of transcript NM_000218.2, and there are 34 labeled P/LP variants in ClinVar between the methionine at codon 1 and the methionine at codon 159. In three other initiation codon variants in the same study (all elevated to PVS1_VeryStrong by ClinGen’s Expert Panel),^12^ the number of ClinVar P/LP variants upstream of the closest potential in-frame start codon was 16, 80, and 41 (Table S2). Give this evidence, we suggest the PVSI strength level of NM_000218.2:c.2T>C(p.Met1?) in *KCNQ1* should be modified from PVS1_Moderate to PVS1_VeryStrong.

The *GAA* gene has 20 exons. Variant NM_000152.4:c.1327-2A>G (p.?) is in the splice acceptor site of exon 9 of this gene. ClinGen’s SVI Working Group classified the variant as PVS1_moderate because the reading frame is preserved, the region’s role is unknown, and the altered protein accounts for less than 10% of the whole protein. However, our assessment found a cryptic splice site in the 16th nucleotide sequence 5’ upstream of exon 9 (Table S3). Adding the 111 nucleotide sequence of exon 9, the total number of nucleotides is 127, which is not divisible by three. We therefore determine that the reading frame is disrupted following frameshift and NMD, and this variant should be classified as PVS1_VeryStrong.

Variant NM_000277.2:c.168+1G>A(p.?) is in the splice donor site of exon 2 of the *PAH* gene. ClinGen and AutoPVS1 agreed that cryptic splice sites were not observed in nearby sequences, leading to exon skipping. However, ClinGen’s SVI Working Group determined a rating of PVS1_Moderate because the function of the skipped exon is unknown.^12^ By contrast, we propose that the region is critical because 25 pathogenic/likely pathogenic variants were found but no benign variants. Thus, the variant should be rated PVS1_Strong.

### Comparative analysis with InterVar

We downloaded InterVar and processed dataset 1 with default settings for comparison. InterVar only produced two outcomes: 1, indicating variants compatible with PVS1 criteria (PVS1_VeryStrong), and 0, indicating variants which do not meet the criteria (Unmet).^19^ As a result, only 52% (29/56) of InterVar’s interpretations were consistent with ClinGen’s results, significantly lower than the concordance of AutoPVS1 (Figure 2). Of 15 variants (5 nonsense, 6 frameshift, 4 initial codons) which InterVar determined did not meet the PVS1 criteria, ClinGen’s Working Group assigned 10 variants a PVS1_VeryStrong rating, 4 variants as PVS1_Moderate, and 1 variant PVS1_Strong (Table S4). By comparison, AutoPVS1’s interpretations were nearly all consistent with ClinGen, with the exception of the three cases discussed above. Notably, InterVar assessed all splicing variants as PVS1_VeryStrong, which is an overestimate in comparison with the results from ClinGen’s Working Group and AutoPVS1. Finally, AutoPVS1 was designed to also process exon-level deletions/duplications, which InterVar currently cannot do.

### Impact of PVS1 interpretation

To analyze the impact of ClinGen’s recommendations on PVS1 interpretation, we extracted a total of 28,586 putative LoF variants from the most prevalent transcripts of 578 genes in ClinVar, including 15,824 frameshift variants, 7,758 nonsense variants, 4,717 splicing variants, and 287 initiation codon variants. Overall, we found that 27.6% of the variants should be assigned a lower final strength level, with 17.4% due to variant-specific issues and 10.2% due to disease mechanism considerations.

Comparing the strength levels in the two steps of the analysis, we find that 82.6% of the variants met the PVS1_VeryStrong criteria in the first step, but the proportion decreased to 72.4% following the second step. Likewise, the proportion of variants that did not merit assignment of a PVS1 strength level increased significantly, from 0.9% in the first step to 10.5% in the second step. These results highlight the importance of considering disease mechanisms in variant interpretations.

Moreover, only 59.3% (2,799/4,717) of splicing variants were assigned a very strong level for pathogenicity in the first step; that is, 40.7% (1,918/4,717) of splicing variants were assigned a lower PVS1 strength level, a significantly greater fraction than in frameshift variants (13.6%) and nonsense variants (10.2%). This is mainly attributed to the fact that 35.4% (1,670/4,717) of splicing variants preserved the reading frame even though an exon was skipped or a cryptic splice site was activated. We therefore suggest that the classification of splicing variants should not rely heavily on LoF contributions and encourage the integration of experimental or case-level evidence to establish their pathogenicity. Our findings reinforce the necessity of considering variant-specific issues in interpretation.

After initiation codon variants were examined by ClinGen’s Expert panels, ClinGen proposed to increase the strength level to very strong strength when ≥1 pathogenic variant(s) upstream of the closest potential start codon was identified.^12^ Based on a more stringent threshold which we pre-defined (see Material and Methods), we still found that 61.4% of variants merited strength levels higher than moderate in the first step, even without weighing disease mechanisms (51.6% in PVS1_VeryStrong and 9.8% in PVS1_Strong, respectively).

### Analysis by clinical significance and allele frequency

Translating the ACMG/AMP criterion system into a quantitative framework, PVS1 was assumed to have an odds of pathogenicity of 350, much higher than other pathogenic criteria (18.7 for strong evidence, 4.3 for moderate evidence, 2.08 for supporting evidence).^35^ That is, a variant with a PVS1_VeryStrong strength level is prone to be pathogenic. Based on this assumption, we attempted to explore the character of variants with different PVS1 strength levels. We stratified variants by clinical significance retrieved from ClinVar^22^ and their allele frequency from the gnomAD database^26^. Unsurprisingly, pathogenic/likely pathogenic variants were more likely to have a PVS1_VeryStrong strength level, significantly more so than benign/likely benign variants (Figure 3a; Fisher’s exact test, *P*<0.001; 76.2% vs 17.6%). Similarly, we detected a strong enrichment of PVS1_VeryStrong in low-frequency variants (*P*<0.001; Figure 3b).

**Figure 3.**
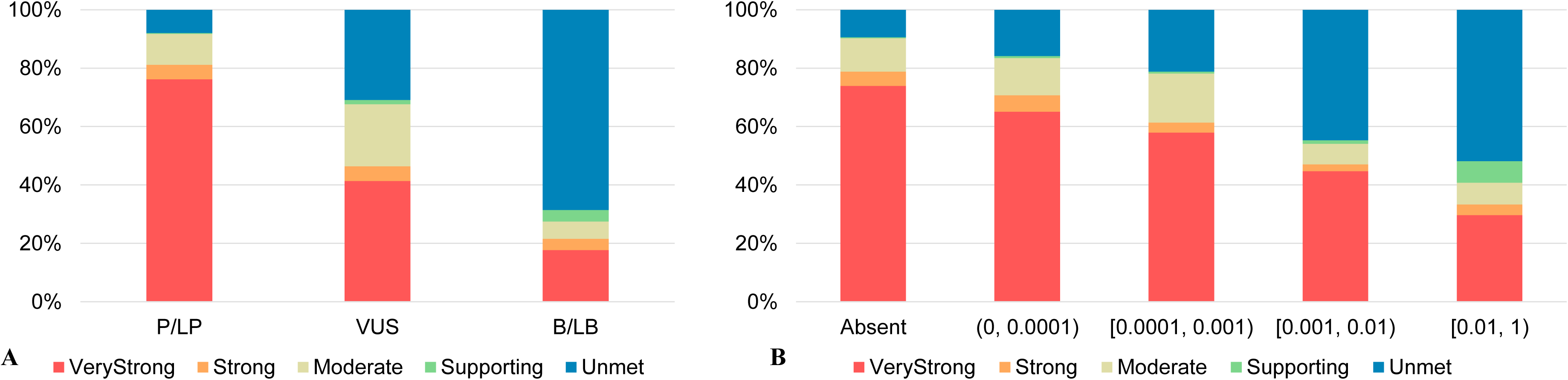
PVS1 strength levels of 30,526 null variants by clinical significance (A) and allele frequency (B). P, pathogenic; LP, likely pathogenic; VUS, variant of uncertain significance; LB, likely benign; B: benign.

### AutoPVS1 web version

To facilitate the use of AutoPVS1 by users unfamiliar with the command line, we developed a web version which can be used to query all types of null variants (Figure 4). Users first search for a variant by HGVS nomenclature or genomic position (Figure 4A). The results are then displayed in three parts: variant information (Figure 4B), preliminary PVS1 strength level according to the PVS1 criteria (Figure 4C), and final adjusted PVS1 strength level weighing disease mechanisms (Figure 4 D).

**Figure 4.**
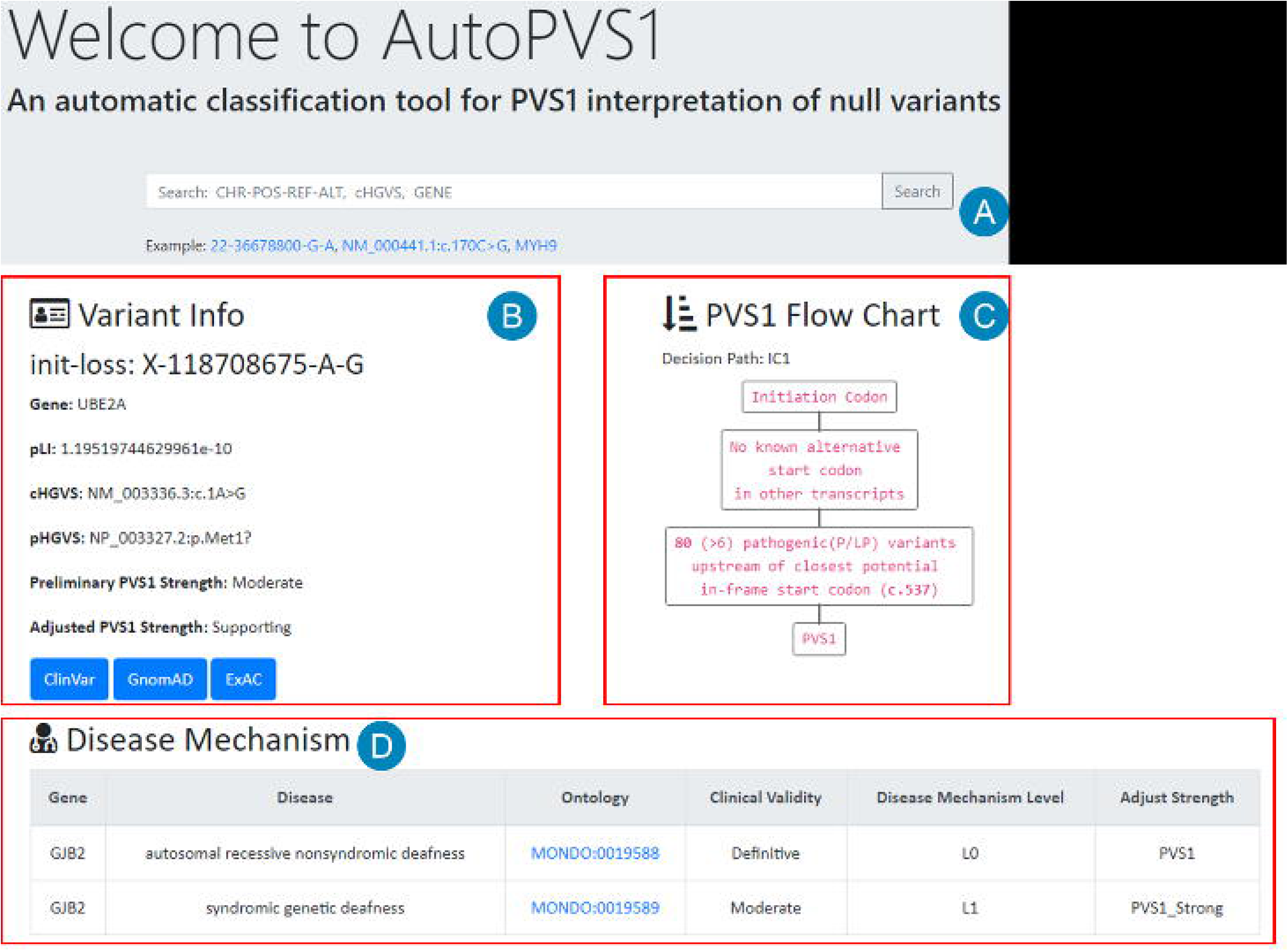
AutoPVS1 web-interface workflow. (A), variant query; (B) variant information; (C), preliminary PVS1 strength levels via variant-specific issues; (D), final adjusted PVS1 strength levels weighing disease mechanisms.

We visualize the decision path for preliminary PVS1 strength levels (Figure 4C). Since a gene may be associated with multiple diseases with different strengths of gene-disease clinical validity, we listed relevant diseases retrieved from the Online Mendelian Inheritance in Man (OMIM) database if the gene was not curated and the clinical validity should by selected by users (Figure 4D). For genes that have been curated by ClinGen, the gene-disease clinical validity is provided.

## Discussion

In this study, we developed a computational tool named AutoPVS1 for PVS1 interpretation based on the recommendations of ClinGen’s SVI Working Group.^12^ The software is available as a command-line program and through a web interface. To the best of our knowledge, this is the first publicly available tool for PVS1 interpretation using the latest PVS1 specifications. Considering the high prevalence of LoF variants in the human genome^10^ and the complexity of PVS1 specifications,^12^ AutoPVS1 meets an urgent need by enabling biocurators to easily assign accurate, reliable and reproducible PVS1 strength levels in the process of variant interpretation.

We would like to emphasize that PVS1 interpretation consists of two independent steps. The first step is to determine preliminary strength levels by addressing issues specific to each variant type. The second step, which involves consideration of disease mechanisms, should be manually adjusted in practice. This is mainly because mutations in the same gene cause a broad range of diseases and the gene-disease associations are heterogeneous, which finally determines whether how preliminary PVS1 strength levels should be altered, if at all.^12^ For example, *GJB3* gene (MIM: 603324) had a “Strong” association with erythrokeratodermia variabilis, whereas it has a “Disputed” association with nonsyndromic genetic deafness.^36^ Therefore, PVS1 should not be applied at any strength level when interpreting a null variant in *GJB3* gene with hearing loss, but it is applicable for erythrokeratodermia variabilis.

Of the ~60,000 putative LoF variants in ClinVar, we included only 28,586 variants in our analysis because their gene-disease clinical validities are freely available from ClinGen. Nevertheless, the preliminary strength levels of all 60,000 variants have been pre-annotated and embedded in the tool. We will continue to update the gene/disease list over time. In addition to ClinGen’s efforts, the development of AutoPVS1 relied on a number of open-access resources such as ClinVar,^22^ RefSeq,^23^ UniPort,^25^ highlighting the importance of data sharing through centralized databases.^37^

One of the advantages of AutoPVS1 is its comprehensive consideration of the specifications for different variant types. This led to a high interpretation concordance (95%) between AutoPVS1 and ClinGen’s SVI Working Group in our benchmarking assessment. A further analysis of 28,586 LoF variants by AutoPVS1 demonstrated that 27.6% of variants should be assigned decreased strength levels, consisting of 17.4% based on variant-specific issues and 10.2% based on disease mechanism considerations. The results indicate the great impact of ClinGen’s PVS1 interpretation recommendations in interpreting LoF variants.

One important caveat we would like to stress is the impact of transcriptional architecture on PVS1 interpretation. Transcript selection is crucial for all null variant types because variants can have different consequences for each transcript. Although we pre-defined the most biologically-relevant transcript for each gene in AutoPVS1, it should be noted that multiple biological transcript isoforms may exist for one gene.^38^ In this circumstance, curators need to carefully review the expression and function of each transcript.

In addition, users should note that as AutoPVS1 is a generic computational tool to interpret null variants, certain thresholds are determined empirically, and it may be more appropriate to modify them according to the characters of specific diseases. For example, we determined that a null variant with an allele frequency greater than 0.1% should be used to support the insignificance of the exon. In comparison, 0.3% was used for hearing loss transcript curation.^21^ The threshold issue can be addressed through adjustable parameters by command-line users.

AutoPVS1 also has limitations. When considering disease mechanisms, it is recommended to count the number of pathogenic LoF variants based on a pathogenicity evaluation without PVS1, and 10% of those LoF variants should be associated with the phenotype.^12^ However, we extracted and counted pathogenic LoF variants from ClinVar directly without considering the pathogenicity without PVS1 and the phenotype. We made this compromise because no such database is currently available, although it may lead to bias in our analysis. However, we believe this will be a minor issue for AutoPVS1 in practice because we also provide users with an alternative mechanism to evaluate the importance of LoF of these genes. To minimize the impact of any bias introduced by our LoF variant count, we present the pLI of each gene from ExAC^26^ as a reference in the web version of AutoPVS1. pLI is a probability that a gene is intolerant to a LoF mutation. Genes with a pLI greater than 0.9 are regarded as extremely intolerant of loss of function.^26^

ClinGen’s SVI Working Group elaborated the recommendations for PVS1 interpretation, which have great impact on variant interpretation. While the elaborations may improve the consistency of interpretation, they also increase the complexity of analysis and the bioinformatics requirements. In this study, we developed AutoPVS1 to streamline the assignment of PVS1 strength levels. We also developed a user-friendly interface for non-command-line users. Our results demonstrate that AutoPVS1 is an accurate, reproducible, and reliable tool, which greatly facilitates clinical PVS1 interpretation and will therefore be of importance to curators.

## Supplemental Data

Supplemental Data include 1 figure and 4 tables.

## Acknowledgments

Not applicable.

## Declaration of Interests

Jiale Xiang, Jiguang Peng, and Zhiyu Peng were employed at BGI at the time of submission. No other conflicts relevant to this study should be reported.

## Web Resources

ClinVar, https://www.ncbi.nlm.nih.gov/clinvar/

The Exome Aggregation Consortium (ExAC): http://exac.broadinstitute.org/

The Genome Aggregation Database (gnomAD): https://gnomad.broadinstitute.org/

Online Mendelian Inheritance in Man (OMIM): https://omim.org/

RefSeqGene/the Locus Reference Genomic collaborations: https://www.lrg-sequence.org

The Reference Sequence (RefSeq): https://www.ncbi.nlm.nih.gov/refseq/

UCSC Genome Browser: https://genome.ucsc.edu/cgi-bin/hgGateway

The International Mouse Phenotyping Consortium (IMPC): http://www.mousephenotype.org/

wInterVar: http://wintervar.wglab.org/

Web version of AutoPVS1:

